# Modeling the metabolic evolution of mixotrophic phytoplankton in response to rising ocean surface temperatures

**DOI:** 10.1101/2022.07.21.501028

**Authors:** Logan M. Gonzalez, Stephen R. Proulx, Holly V. Moeller

## Abstract

**Background:** Climate change is expected to lead to warming in ocean surface temperatures which will have unequal effects on the rates of photosynthesis and heterotrophy. As a result of this changing metabolic landscape, directional phenotypic evolution will occur, with implications that cascade up to the ecosystem level. While mixotrophic phytoplankton, organisms that combine photosynthesis and heterotrophy to meet their energetic and nutritional needs, are expected to become more heterotrophic with warmer temperatures due to heterotrophy increasing at a faster rate than photosynthesis, it is unclear how evolution will influence how these organisms respond to warmer temperatures. In this study, we used adaptive dynamics to model the consequences of temperature-mediated increases in metabolic rates for the evolution of mixotrophic phytoplankton, focusing specifically on phagotrophic mixotrophs.

**Results:** We find that mixotrophs tend to evolve to become more reliant on phagotrophy as temperatures rise, leading to reduced prey abundance through higher grazing rates. However, if prey abundance becomes too low, evolution favors greater reliance on photosynthesis. These responses depend upon the trade-off that mixotrophs experience between investing in photosynthesis and phagotrophy. Mixotrophs with a convex trade-off maintain mixotrophy over the greatest range of temperatures; evolution in these “generalist” mixotrophs was found to exacerbate carbon cycle impacts, with evolving mixotrophs exhibiting increased sensitivity to rising temperature.

**Conclusions:** Our results show that mixotrophs may respond more strongly to climate change than predicted by phenotypic plasticity alone due to evolutionary shifts in metabolic investment. However, the type of metabolic trade-off experienced by mixotrophs as well as ecological feedback on prey abundance may ultimately limit the extent of evolutionary change along the heterotrophy-phototrophy spectrum.

## BACKGROUND

Anthropogenic climate change is expected to increase oceanic surface temperatures [1], with cascading effects on ocean ecosystems. An important outcome of warmer temperatures will be faster metabolic rates due to the temperature sensitivity of chemical reactions [2,3] This thermal acceleration will influence carbon cycling processes that depend on metabolic rates, including ecosystem respiration and gross primary production [4]. Since respiration responds more strongly to temperature than photosynthesis [5,6] carbon cycling in some regions may shift to favor increased CO_2_ production, driving a positive feedback loop with implications for the future of the biosphere and Earth’s climate [7].

In addition to the direct effects of temperature on their metabolic rates, organisms, especially microorganisms with their relatively fast generation times, will likely evolve in response to warmer temperatures. These evolutionary responses may either exacerbate or mitigate short-term plastic responses [8]. For example, phytoplankton, primary producers which account for nearly half the planet’s photosynthetic net primary production [9], may evolve by shifting their optimal growth temperature [10] or decreasing rates of both photosynthesis and respiration to compensate for temperature-dependent changes in metabolism [11]. A better understanding of microbial evolutionary adaptations to warming, and their consequences for the carbon cycle, will likely increase the quality of long-term climate forecasts.

Today, many microbial primary producers are known to be mixotrophic [12]: they combine photosynthesis and heterotrophy to meet their energetic and nutritional needs. Thus, these lineages may act as carbon sinks (via photosynthesis) or sources (via heterotrophy) depending on the environmental context. Mixotrophs are increasingly recognized as important components of surface ocean ecosystems, where they may dominate bacterivory [13] and regulate carbon export via the biological pump [14]. Because mixotrophs can continue to obtain nutrients through heterotrophy when inorganic nutrient concentrations are low, they are expected to dominate late-season and oligotrophic ecosystems [15]. These nutrient-limited, mixotroph-favoring conditions are expected to become more widespread in a warmer, more stratified ocean [16], so accurate predictions of mixotroph responses to warming temperatures are needed. However, mixotrophs’ dual metabolism complicates predicted plastic and evolutionary responses because both sets of metabolic processes are to some extent competing for finite cellular resources and differ in their temperature dependencies.

Here, we focus on constitutive mixotrophs, single-celled planktonic eukaryotes that contain chloroplasts yet retain the capacity for feeding, typically on bacteria [15,17]. Thus, our mixotrophs represent a specific subset of “mixoplankton,” planktonic protists that obtain nourishment from photosynthesis, phagotrophy, and osmotrophy [18] The different thermal scaling of heterotrophy and photosynthesis [5] has led to the prediction that these mixotrophs (like ecosystems) should become more heterotrophic at warmer temperatures [19]. This prediction has mixed empirical support however: In short-term studies of phenotypic plasticity, freshwater mixotrophs from the genus *Ochromonas*, marine *Isochrysis galbana*, and freshwater *Chromulina* sp. have shown increased rates of bacterivory with temperature and overall shifts towards a more heterotrophic metabolism [19-21]. In contrast, the freshwater dinoflagellate *Dinobryon sociale* and marine dinoflagellate *Karlodinium armiger* show increased relative contributions of photosynthesis with temperature [22-23] indicating that mixotrophs’ underlying physiological constraints will shape their thermal response [22] Further, it is unclear how mixotrophs will respond over evolutionary timescales due to the costs they experience from maintaining two fundamentally different forms of metabolism [24-26].

Eco-evolutionary theory, which accounts for the interplay between evolutionary and ecological processes [27,28], can be a useful framework for understanding the evolution of organisms in the context of their environments. Previous work applying eco-evolutionary theory to model mixotroph evolution has shed light on the factors contributing to metabolic specialization in mixotroph lineages by focusing on cell size [29] and the costs and benefits of heterotrophic and photosynthetic metabolism [30]. We applied adaptive dynamics [31,32] to explore how eco-evolutionary feedback influences mixotroph metabolism in response to temperature, assuming a trade-off exists between investment in photosynthesis and phagotrophy. Previous studies of resource trade-offs have shown that evolutionary outcomes are largely dependent on the set of attainable phenotypes, or fitness set [33,34], which is a function of physiological constraints on resource acquisition as well as interactions between resources themselves. Resources can be classified as complementary, acting in a cooperative manner; antagonistic, inhibiting acquisition of the alternative resource; or perfectly substitutible, in which equilibrium can be maintained by substituting the intake of one resource by a proportional intake of the alternative resource [35-37]. Trade-offs resulting from resource acquisition characteristics may be convex, tending to favor generalists that acquire both resources in similar quantities, concave, penalizing intermediate strategies and producing the highest growth rates when specializing on a single resource, or linear, which favor neither specialist or generalist strategies [34, 36-38]. In addition, evolutionary outcomes can become more complex in the presence of frequency-dependent selection [38-39].

Our model predicts that mixotrophs may evolve to become more phagotrophic, and therefore more dependent on heterotrophy, with rising temperatures. However, the evolved investment strategy is influenced by both the type of trade-off experienced between photosynthesis and phagotrophy and ecological feedbacks on prey density, which together influence the contribution of photosynthesis and phagotrophy to mixotroph growth

## METHODS

We model the interaction between a mixotroph *M* and its bacterial prey *B* living in a well-mixed water column. An individual mixotroph grows through photosynthesis P(*θ*, z, I_in_, T, *M*) and grazing G(*θ*, T, *B*), both of which depend on temperature T, and has a constant *per capita* mortality rate l. Thus, the change in mixotroph population density over time is given by:

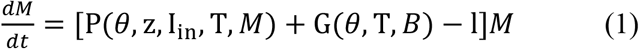

Both photosynthetic rates and grazing rates are dependent on a combination of environmental factors—e.g., mixotroph (*M*) and bacteria (*B*) density, temperature (T), and input light (I_in_)—and physiological factors—e.g., mixotroph investment in phagotrophy (*θ*) and the trade-off parameter (z) (Figure 1, Table 1).

**Table 1.**
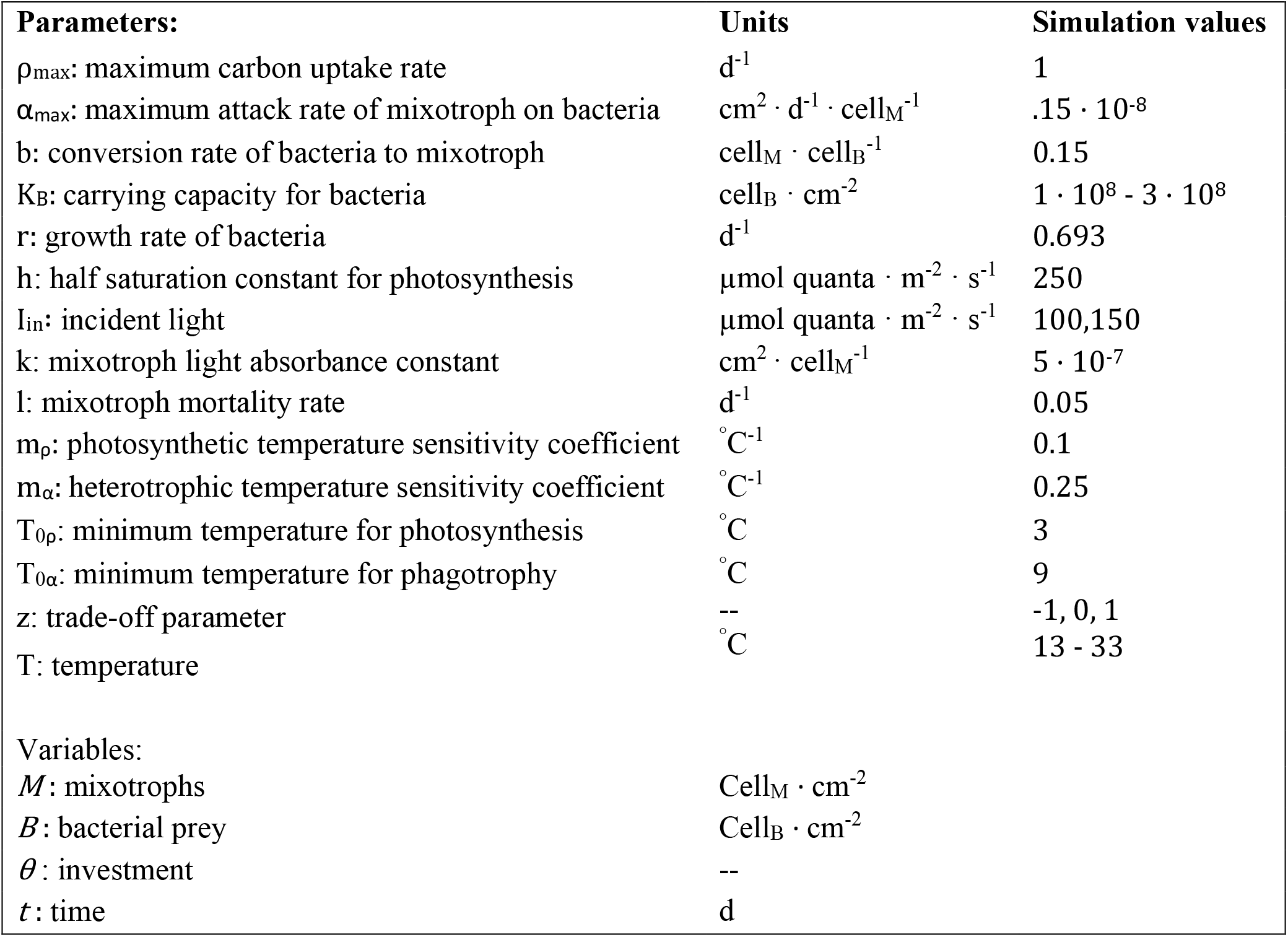
Model parameters with units and values used in simulations.Simulation values for mixotroph parameters were chosen based on the range of values used in Moeller et al. 2019. Bacterial prey parameters, r and K_B_, were selected that clearly demonstrated the general qualitative outcomes of each trade-off type over the simulated temperature range.

**Figure 1.**
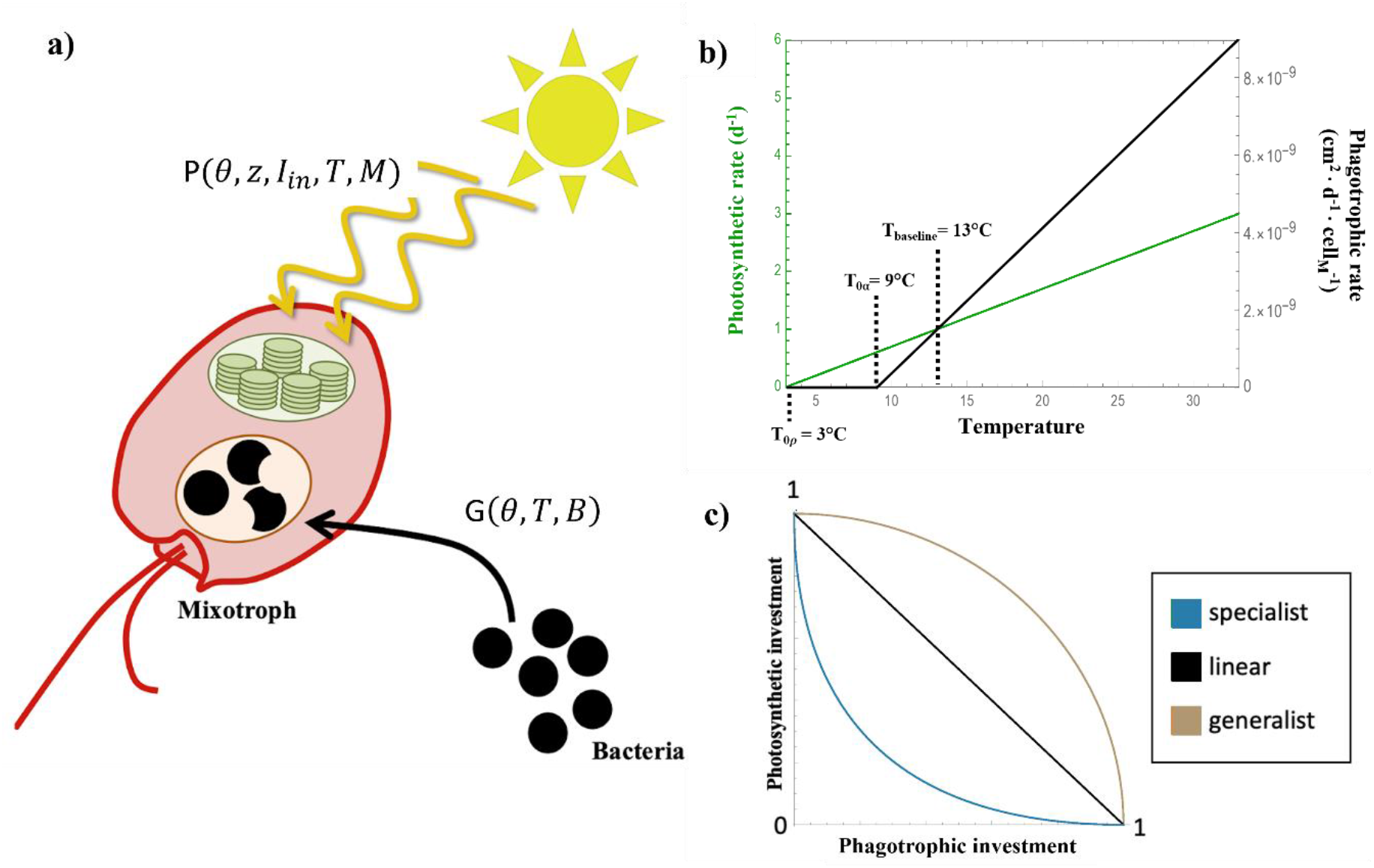
Diagram of model components. (a) Mixotrophs grow by combining photosynthesis (harvest of sunlight, yellow lines, using a chloroplast, green oval) and phagotrophy (consumption of bacteria, black circles). The rates of both photosynthesis and phagotrophy depend upon temperature T and phagotrophic investment *θ*. (b) Both photosynthesis and phagotrophy depend on temperature. We assume that, over the temperature range of interest, thermal responses can be approximated as linear, with phagotrophy (black line) more sensitive to increases in temperature than photosynthesis (green line). (c) Mixotroph investment in photosynthesis and phagotrophy is constrained by a trade-off: Assuming that metabolic resources are finite, a mixotroph cannot simultaneously achieve maximum rates of both forms of metabolism. We model three different types of trade-offs: convex (light brown), which produces generalist-type mixotrophs that tend to maintain both forms of metabolism; concave (blue), which produces specialist-type mixotrophs that invest fully in either photosynthesis or phagotrophy; and linear (black).

Photosynthetic growth follows the model developed by Huisman & Weissing (1994) [40]. Huisman and Weissing’s model begins with the assumption that photosynthesis is a saturating function of light. The photosynthetic rate at a given irradiance ρ(I) is a function of the maximum photosynthetic rate ρ_max_, the irradiance I, and the light level at which photosynthesis reaches half of its maximum rate h:

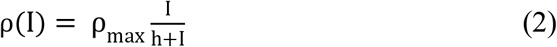

For a population of mixotrophs *M* living in a well-mixed water column, the light environment I is a function of depth. In particular, light is assumed to attenuate from a surface input irradiance of I_in_ with depth due to absorption by photosynthetic cells which have a per-cell absorptivity k. Huisman and Weissing computed the average photosynthetic rate across depth, and obtained the *per capita* photosynthetic growth rate, defined here as P(*θ*, z, I_in_, T, *M*):

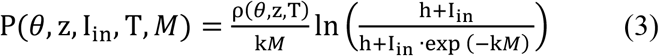

Mixotrophs also obtain resources by grazing on bacteria. Therefore, we included a grazing term (sensu Moeller et al. 2019 [41]) assuming that, at relevant bacterial population densities, mixotroph grazing on bacteria at density *B* can be approximated using a Type I functional response [42], where *α*(*θ*, T) represents the mixotroph attack rate and b represents the conversion efficiency of captured bacteria into mixotroph biomass:

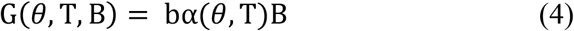

Metabolic rates are known to scale with temperature [3]. To simplify our model, we assume that photosynthetic and grazing rates are linear functions of temperature (Figure 1B):

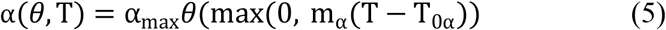

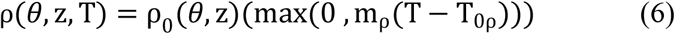

Here, *θ* is an investment parameter that controls the reliance of mixotrophs on phagotrophy and photosynthesis. When *θ* = 0, mixotrophs invest solely in photosynthesis. When *θ* = 1, mixotrophs invest solely in phagotrophy. T_0*α*_ and T_0ρ_ represent the temperatures at which the maximum rates of phagotrophy and photosynthesis reach zero, respectively. The temperature coefficients mρ and m*α* were estimated by calculating the Q10-values from the activation energies of carbon fixation and grazing previously measured for *Ochromonas* sp. [19,43]. Although thermal reaction norms are typically unimodal nonlinear functions of temperature [44], in our case assuming a linear function allows us to avoid introducing an underlying thermal nonlinearity into our system. This allows us to attribute any modeled non-linear evolutionary responses to ecological feedback rather than underlying assumptions about the shape of the thermal response. The use of an exponential function did not lead to qualitatively different results (Additional File 1: Figure S1). For similar reasons, we chose to fix bacterial growth rates rather than making them increasing functions of temperature. Again, this simplification allows us to isolate the role of mixotroph thermal responses in driving evolution.

We introduced a trade-off between photosynthesis and phagotrophy by allowing the photosynthetic rate to vary as a function of phagotrophic investment *θ* and a shape parameter z.

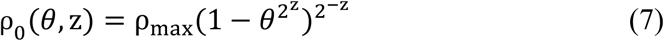

The shape parameter z determines the curvature of the photosynthesis-phagotrophy trade-off strategy for mixotrophs (Figure 1C). Mixotrophs with shape parameters of z < 0 are defined as specialists because investment in one metabolic strategy comes at a large cost in investment in the alternative. In contrast, mixotrophs with shape parameters of z > 0 are defined as generalists because they are capable of maintaining relatively high levels of photosynthesis and phagotrophy simultaneously. When z = 0, there is a linear trade-off between photosynthesis and phagotrophy. For simplicity, we limited our analysis to z = 1, z = -1, and z = 0.

Assembling these components, mixotroph population dynamics are governed by the equation:

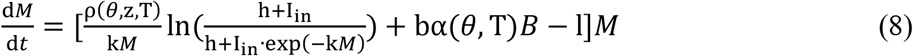

Bacterial prey populations are assumed to grow logistically, with a growth rate r and carrying capacity K_*B*_, and are grazed on by mixotrophs *M*:

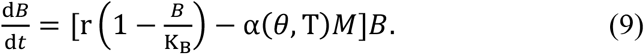

Both mixotroph and bacterial abundances are measured in cells per area (see Table 1), because, following Huisman & Weissing (1994), we are using depth-integrated measurements of well-mixed populations.

## ANALYSIS AND RESULTS

We ran model analyses and simulations using Mathematica version 12. Data and code are available at https://github.com/lg11235/mixotroph-adaptive-dynamics. (Note to reviewers: We will update this to a permanent DOI upon manuscript acceptance.) We parameterized our model following previously collected data on marine *Ochromonas* strains [41], which are constitutive mixotrophs representative of those we are modeling. Temperature sensitivities were parameterized following data reported in Wilken et al. (2013) [19].

We applied an adaptive dynamics framework to our ecological model to track the evolution of mixotroph growth strategies (phagotrophic and photosynthetic investment) as a function of temperature (Additional file 1: Section S2). In effect, we assume that ecological dynamics are based on the resident mixotroph allele and come to equilibrium before new mutations altering the growth strategy arise. When a new mixotroph allele appears, the mutant allele may be lost when rare or begin to spread in the population. If a mutant allele spreads when rare, it will continue to fixation unless frequency dependence causes the relative fitness of the resident allele to increase above that of the mutant allele. When ecological dynamics occur on a fast time-scale relative to mutation, the evolutionary trajectory can be modeled as a series of replacement events [45-47], So long as we consider small changes in the mixotroph strategy, if a mutant invades the resident and the resident cannot invade the mutant, then the mutant will replace the resident [48-49]. If we observe mutual invasibility, on the other hand, that is sufficient to show that neither allele represents an evolutionary stable point, but not sufficient to show that coexistence or adaptive dynamic branching will occur. Based on these considerations, we model the evolution of mixotroph strategies as a trait-substitution sequence and note that alternative formulations of evolution within the model are likely to generate the same conclusion [50].

Mutant mixotrophs with investment strategy *θ*_*mut*_ arise by mutation as rare genotypes within the resident mixotroph population with investment strategy *θ*_*res*_. The mutant growth rate is dependent on the equilibrium densities of the prey and resident mixotroph, giving the following *per capita* growth rate equation for a mutant, defined as the invasion fitness [31]:

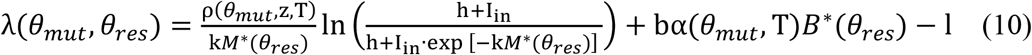

where *M*^*^(*θ*_*res*_) and *B*^*^(*θ*_*res*_) represent the population densities of resident mixotrophs and bacteria at equilibrium. When *λ*(*θ*_*mut*_, *θ*_*res*_) > 0 and *λ*(*θ*_*res*_, *θ*_*mut*_) < 0 (fitness of a single resident invading a population of mutants at equilibrium) the mutant invades and replaces the resident mixotroph population. This invasion-replacement process occurs until an investment strategy that is unbeatable is reached (*θ*_*ESS*_), resulting in an evolutionarily stable state (ESS) where the system is locally stable to further invasion. We used our model to predict the evolutionarily stable phagotrophic investment strategies for mixotroph as a function of temperature, *θ*_*ESS*_(T). Since *M*^*^(*θ*_*res*_) and *B*^*^(*θ*_*res*_) do not have closed-form solutions, *θ*_*ESS*_(T) was solved numerically. We considered a temperature range from 9^°^C to 33^°^C at varying light levels and maximum prey carrying capacities.

### Assessing mixotroph invasibility across heterotrophic investment space

Analyzing invasion fitness for all combinations of *θ*_*mut*_ and *θ*_*res*_ and identifying the regions where *λ*(*θ*_*mut*_, *θ*_*res*_) > 0 using pairwise invasibility plots (PIPs) provides a useful way to visualize the global evolutionary behavior of the model [31]. PIPs were plotted for mixotrophs with specialist, linear, and generalist trade-offs at 13^°^C, 18^°^C, and 23^°^C (Figure 2).

**Figure 1.**
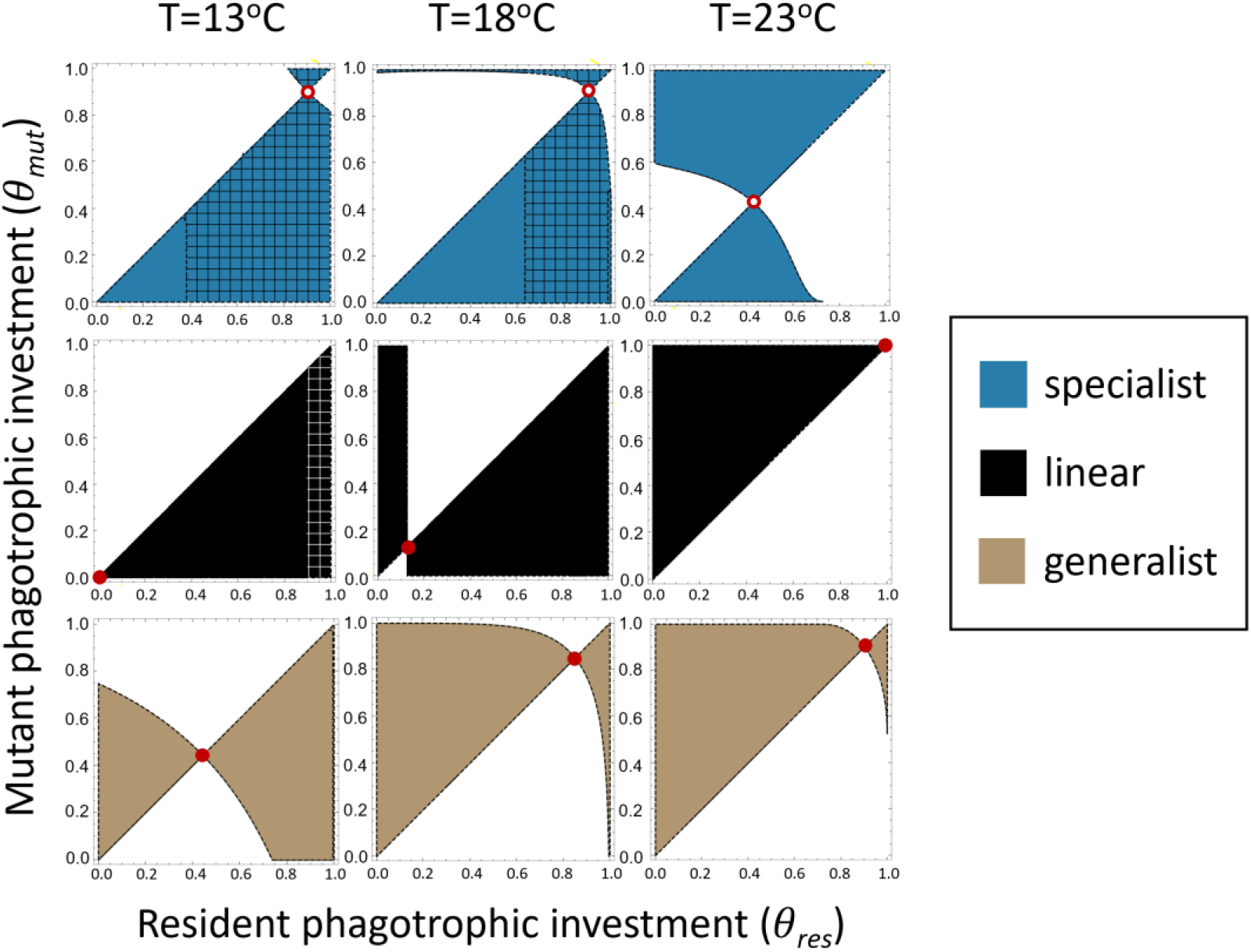
Pairwise invasibility plots for mixotrophs of varied trade-off functions. Shaded areas indicate the mutant strategies that can successfully invade residents with a particular strategy indicated on the x-axis. Evolutionary singularities are depicted by open and closed circles to indicate evolutionary stability or instability, respectively. Gridlines indicate regions of trait space in which resident mixotrophs are nonviable and mutants invade an empty environment. Top row: Specialist mixotrophs (z = -1) evolve to become completely specialized (i.e., *θ*_*ESS*_ = 0 or *θ*_*ESS*_ = 1) depending on the starting metabolic investment) regardless of temperature. Middle row: Mixotrophs with linear trade-offs (z = 0) evolve to invest more in phagotrophy as temperatures increase. Bottom row: Generalist mixotrophs (z = 1) are functionally mixotrophic (0 < *θ*_*ESS*_ < 1) across a wider range of temperatures compared to mixotrophs with linear trade-offs. Evolutionary branching was not found to occur for specialist mixotrophs over this temperature range. K_B_ = 1 · 10^8^ cell_B_ · cm^-2^ and I_in_ = 100 µmol quanta · m^-2^ _· s_^-1^ used to generate plots.

In some cases, particularly for specialist trade-offs, resident mixotroph population sizes were negative and mixotroph population sizes were set to 0. Thus, mutant mixotrophs in these regions invade an empty environment. For specialist mixotrophs, this caused *θ*_*ESS*_ = 1 to be a nonviable evolutionary strategy at low temperatures. Across all trade-off types, evolution to *θ*_*ESS*_ produced equilibrium mixotroph densities that were lower than the maximum achievable density (Additional file 1: Section S3).

For specialist mixotrophs, the evolutionarily singular strategy was a repeller point under most simulated conditions, leading to extreme investment in either photosynthesis or phagotrophy depending on the investment strategy of the initial resident mixotroph. Specialist mixotrophs were unable to converge on intermediate *θ*_*ESS*_ values between 0 and 1 while the sole evolutionarily singular strategy was a repeller point. At extreme temperatures, an additional evolutionarily singular strategy arises for specialist mixotrophs that is always a branching point. For values of 0 > z > -1, branching is possible at more moderate temperatures due to reduced trade-off strength (Additional file 1: Section S4).

*θ*_*ESS*_ for linear trade-off mixotrophs varied with temperature, converging on values at or near total photosynthetic specialization at low temperature but increasing towards higher phagotrophic investment at higher temperatures. For parameters in which linear trade-off mixotrophs converged towards an intermediate *θ*_*ESS*_, the fitness landscape was entirely flat. This means that at *θ*_*ESS*_, the invasion fitness of all mutants with an alternative strategy is zero [36]. However, since this result is only possible in the case of linear trade-offs, we did not explore whether or not this results in coexistence of an arbitrary number of alternative investment strategies or eventual convergence toward the original resident strategy.

The trend for generalist mixotrophs was similar to that of linear trade-off mixotrophs, but favored an intermediate *θ*_*ESS*_ across a larger range of temperatures. This reduced sensitivity of *θ*_*ESS*_ to temperature suggests that generalist mixotrophs experience less selective pressure with respect to metabolic strategy and are able to remain functionally mixotrophic with changing temperature more easily than mixotrophs with linear or specialist trade-offs.

### θ_ESS_ varies with temperature and prey availability

We computed *θ*_*ESS*_ (T) as a function of the light and prey resource landscape (Figure 3).

**Figure 3.**
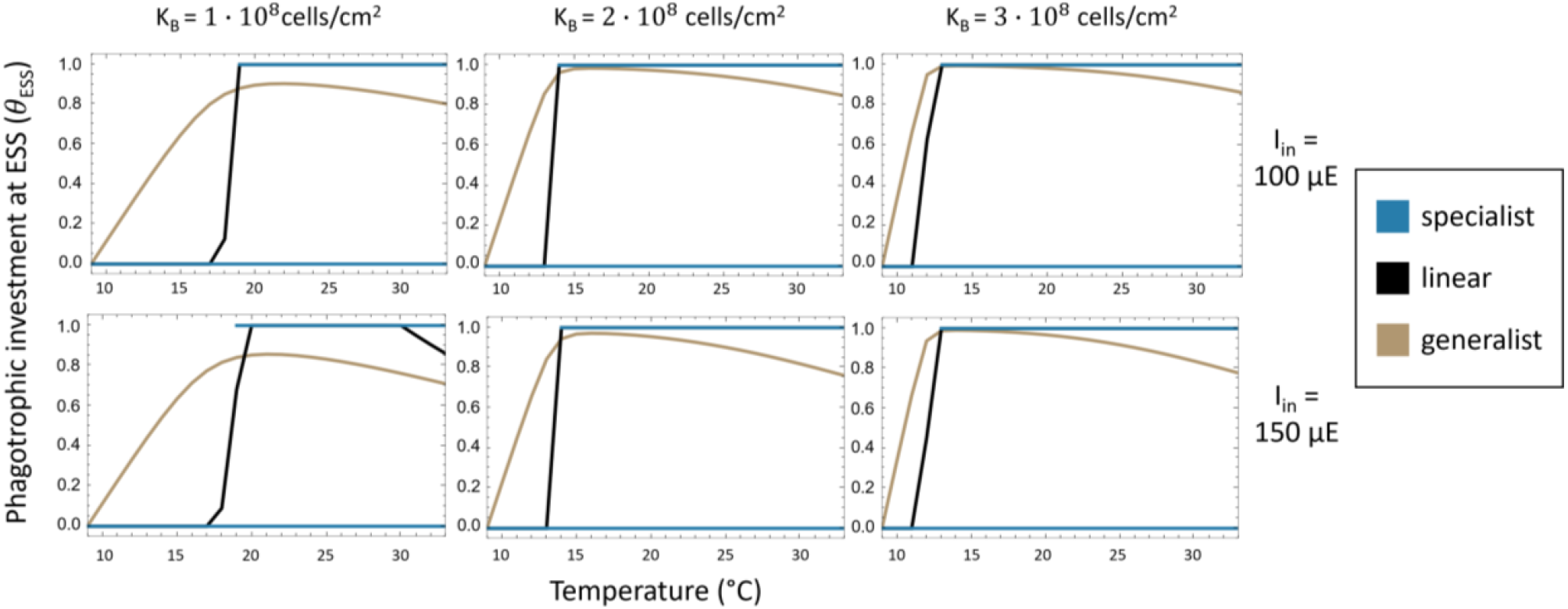
*θ*_*ESS*_ as a function of temperature, light, and prey carrying capacity for specialist (blue), generalist (light brown), and linear (black) trade-off mixotrophs. Specialist mixotrophs exhibited evolutionary bistability at *θ*_*ESS*_ = 0 and *θ*_*ESS*_ = 1 across the temperature range simulated however mixotrophs population sizes were negative in some cases at low temperatures. The line at *θ*_*ESS*_ = 1 only shows values for which mixotroph population sizes are positive.

In the case of linear and generalist trade-off mixotrophs, *θ*_*ESS*_ initially increases with rising temperatures. However, at much higher temperatures, *θ*_*ESS*_ begins to decrease, indicating that photosynthetic investment eventually returns to favorability (Additional file 1: Section S5). *θ*_*ESS*_ for mixotrophs with a specialist trade-off function was fixed at *θ*_*ESS*_ = 0 (fully photosynthetic) or *θ*_*ESS*_ = 1 (fully heterotrophic), with *θ*_*ESS*_ = 1 initially being a nonviable evolutionary strategy until a threshold temperature is reached.. This is consistent with the concavity of the specialist trade-off function, which penalizes investment in more than one form of metabolism. Specialist trade-offs were found to be the only trade-off type that lead to evolutionary branching (Additional file 1: Section S4), in which evolution converges towards an evolutionarily singular strategy before diverging into coexisting lineages that reach different evolutionarily stable states [31]. Increasing prey carrying capacity pushed mixotrophs with generalist and linear trade-offs towards higher levels of phagotrophy while increased light availability caused mixotrophs with linear and generalist trade-offs to rely more on photosynthesis.

### Mixotrophic contribution to carbon cycling with rising temperatures

In order to gain insight into how the evolution of mixotroph strategies in response to rising temperatures would influence carbon cycling, we compared model behavior for genetically static mixotrophs (*θ*_*ESS*_ fixed at its baseline value from 13^°^C) to model behavior for evolving mixotrophs *θ*_*ESS*_ allowed to vary with temperature). This allowed us to contrast strictly thermal responses of mixotrophs with their coupled thermal and evolutionary responses. Specifically, we analyzed differences in both mixotroph and bacterial population densities as well as population-level growth rate components (growth contributions from photosynthesis P(*θ*, z, I_in_, T, *M*) · *M* and phagotrophy G(*θ*, T, *B*) · *M*) for mixotrophs at equilibrium (Figure 4). These population level growth rate components were used as proxies for mixotroph carbon fixation and remineralization. Because mixotrophs with a specialist trade-off function (z = -1) exhibit two ESS values over a subset of this range of temperatures, we explored the cases *θ*_*ESS*_ = 0 and *θ*_*ESS*_ = 1 for this trade-off type.

**Figure 4.**
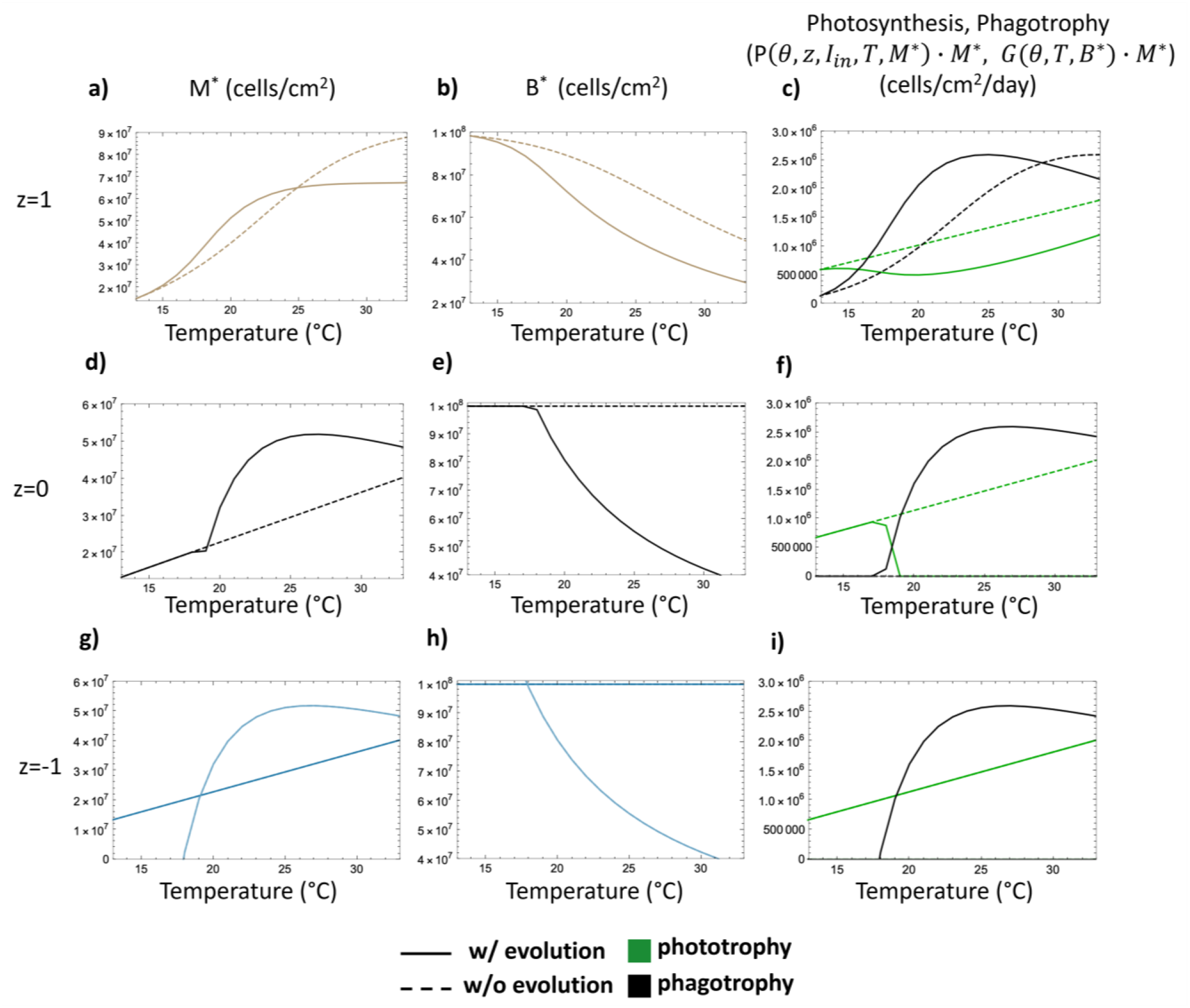
Comparing results of mixotroph-bacterial ecosystems involving either evolving (solid) or unevolving (dashed) mixotrophs as a function of temperature for generalist trade-off (top row) linear trade-off (middle row), and specialist trade-off (bottom row) mixotrophs. Column 1: Mixotroph population densities at equilibrium (*M*\*) vs temperature. Column 2: Bacterial prey population densities at equilibrium (*B\**) vs temperature. Column 3: Photosynthetic and phagotrophic mixotrophic population growth rate components at equilibrium, P(θ, z, I_In_, T, *M\**) · *M\** and G(θ, T, *M\**) · *M\** vs temperature. For plots g and h, outputs for both photosynthetic specialists, *θ*_*ESS*_ = 0, (dark blue) and phagotrophic specialists, *θ*_*ESS*_ = 1, (light blue) are shown. K_B_ = 1·10^8^ cell_B_ · cm^-2^ and I_in_ = 100 µmol quanta · m^-2^ · s^-1^ used to generate plots.

Mixotroph trade-off type determined impacts on carbon cycling (Figure 4). Population sizes of all mixotrophs increased with temperature, although for phagotrophic specialist and linear trade-off mixotrophs (Figure 4d, 4g) population sizes declined at temperatures > 25°C due to sharp declines in bacterial food supply (Figure 4e, 4h). Evolved generalist mixotrophs initially had higher population sizes than their genetically static counterparts, however this trend reversed at higher temperaturesFigure 4a, Additional File 1, Section S5), indicating that evolutionarily stable strategies are not always biomass-maximizing ones. Except when mixotrophs did not invest in phagotrophy, bacterial populations declined with temperature due to accelerating grazing rates and (in most cases) increases in mixotroph abundance.

Generalist mixotroph contributions to ecosystem carbon fixation and remineralization were most complex. Overall, evolution reduced generalist mixotroph contributions to community primary production (Figure 4c). Below 29°C, evolved mixotroph populations contributed more to carbon remineralization through grazing than unevolved lineages (Figure 4c), but at higher temperatures reduced bacterial population density (which feeds back to reduce mixotroph investment in phagotrophy; Figure 3) constrain mixotroph phagotrophy. Thus, at high temperatures, evolved generalist mixotrophs actually contributed less to phagotrophy. In contrast, mixotrophs with linear trade-off functions had increasing population sizes with temperature that initially did not vary with evolutionary history, however there was a large shift at 19°C when *θ*_*ESS*_ = 1 (Figure 4d). This is because, at the reference temperature of 13°C, the linear mixotroph’s *θ*_*ESS*_ was 0 (fully photosynthetic). For this mixotroph type, the selection gradient dictates that, at *θ*_*ESS*_, the mixotroph’s equilibrium population size is the same as a mixotroph with *θ* = 0 unless *θ* = 1 (Additional file 1: Section S6). However, bacterial populations decreased sharply when these mixotrophs could mount an evolutionary response (Figure 4e) due to increased rates of phagotrophy above 18°C (Figure 4f). Because of the trade-off between investments in phagotrophy and photosynthesis, photosynthetic contributions decreased (Figure 4f). For mixotrophs with specialist trade-off functions, *θ*_*ESS*_ did not change with temperature, so evolution did not impact ecosystem-level dynamics. However, due to negative population sizes, *θ*_*ESS*_ = 1 was not a feasible strategy at temperatures lower than 18°C (Figure 4g)

## DISCUSSION

The evolutionary response of mixotrophs to rising temperatures has implications for both the survival and abundance of mixoplankton, and for the cycling of carbon through aquatic ecosystems. The rate of aerobic respiration typically responds stronger to temperature than the rate of photosynthesis [5-6], leading to the hypothesis that mixotrophs should be more heterotrophic at warmer temperatures. Our eco-evolutionary models indicate that over sufficiently long timescales, evolution may compound this prediction, with mixotrophs relying increasingly heavily on phagotrophy as oceanic temperatures rise. However, our findings are complicated by the interplay of physiology (e.g., internal trade-offs between energetic investment in phagotrophy and photosynthesis) and ecology (e.g., the availability of bacterial prey and extent of self-shading) with evolution. In this, our model’s predictions are broadly consistent with experimental data on planktonic evolution. Organisms typically exhibit some capacity for adaptive response to new conditions, but these responses are modulated by environmental conditions [10, 51]. For example, recent work experimentally evolving marine mixotrophs from the Ochromonas genus found that mixotrophs evolved at hotter temperatures tended to eat more and photosynthesize less than un-evolved lineages [52]). But, evidence for evolution varied by mixotroph lineage and light availability (Lepori-Bui et al., in press), suggesting that underlying evolutionary constraints [22] interact with environmental conditions to modulate thermodynamic predictions [19].

Our model predicts that mixotroph responses to temperature are non-monotonic. As temperature increases, mixotrophs on average increase their reliance on phagotrophy. However, at sufficiently high temperatures, investment in phagotrophy counterintuitively decreases (Figure 3, Figure 4). This is because mixotroph responses are shaped by the dynamic tension between internal cellular metabolism and external ecological factors. As mixotrophs become more heterotrophic, they deplete their supply of bacterial prey (an ecological feedback), reducing the favorability of grazing until it no longer becomes beneficial to increase heterotrophic investment (Additional file 1: Section S5). Thus, the direction of selection shifts towards increased investment in photosynthesis. These findings illustrate the complexity of accounting for evolution in the context of ecological feedback. In the absence of these eco-evolutionary feedback (e.g., if prey never became limiting) and assuming a more realistic nonlinear temperature-metabolic rate relationship used in our model, we might expect more phagotrophy than is actually observed at the high temperatures simulated in our model. For example, allowing bacterial growth rates to accelerate with temperature could increase bacterial availability, causing increases in heterotrophic investment similar to those observed with increases in bacterial carrying capacity (Figure 3). We also did not consider co-evolutionary dynamics, even though bacteria are known to evolve rapidly, including to thermal stress [53]), and co-evolution can complicate predator-prey dynamics [54]

Although we chose to use a linear approximation of the thermal responses of phagotrophy and photosynthesis to temperature, no metabolic reactions can realistically increase indefinitely as a function of temperature. At some point, at a maximum critical temperature, enzymes will denature, and the metabolic process will halt. Indeed, previous studies have indicated that phytoplankton evolution may be constrained at this upper thermal maximum [10]. However, in our model, the use of a linear thermal responses is a good approximation for metabolic rates over relatively small temperature increases and allows us to isolate the ecological feedback which drive nonlinearities in the mixotroph’s evolutionarily stable strategy from nonlinearities imposed by the thermal responses themselves.

Over temperature ranges in which scaling of metabolic rates with temperature can be assumed to be monotonic, our model predicted that evolution initially exacerbates increases in heterotrophic rates with respect to temperature while suppressing increases in photosynthetic rates, suggesting that the evolution of mixotroph metabolic strategies under warming ocean surface scenarios will cause a shift towards greater net phagotrophy beyond that predicted from the temperature dependence of metabolic rates alone. The evolution of increased phagotrophy at the expense of photosynthesis in mixotrophs would have important implications for the future of ocean ecosystems and carbon cycling. A reduced capacity for photosynthesis would lower overall primary production, limiting the level of inorganic carbon that can be fixed and used to build biomass. Higher levels of phagotrophy could add to this effect, increasing respiration and leading to more rapid CO_2_ production as well as a potential reduction in concentration of POC in the water column, which would reduce sinking flux via the biological carbon pump [55]. Such effects may lead to a shift from net sinks to net sources of carbon in ocean ecosystems where mixotrophs play a major role. By increasing the rate of biological carbon dioxide flux to the atmosphere, this could create a positive feedback accelerating warming [7]. However, the interplay between rates of environmental change and evolutionary change will be quite important in determining these responses. If evolution is rapid with respect to environmental change, then our approach, which reports only evolutionary endpoints, is a good approximation for mixotrophic tracking of environmental conditions. However, slower rates of evolution may cause carbon flux outcomes intermediate between our predictions. Furthermore, fluctuating environmental conditions can cause time-varying selection pressures that can lead to the evolution of plasticity rather than fixed strategies [56-57] Indeed, mixotrophy may be more common in variable environments because of alternating selection for photosynthesis and phagotrophy.

Trade-off type had a significant impact on the evolutionary response of mixotrophs to warmer temperatures. In our model, mixotrophs with a generalist trade-off function exhibited mixotrophy (i.e., invested in both photosynthesis and phagotrophy simultaneously, 0 < *θ*_*ESS*_ < 1) over a wider range of temperature than mixotrophs with linear trade-off functions or, especially, specialist trade-off functions. Indeed, consistent with the nomenclature of a specialist trade-off type, these mixotrophs tend to specialize by completely investing in either photosynthesis (*θ*_*ESS*_ = 0) or phagotrophy (*θ*_*ESS*_ = 1). These results are consistent with previous studies on resource trade-offs, which have found evolutionary outcomes to depend on the curvature of the trade-off function [34, 36, 37-39] Generally, as the strength of the trade-off increases, extreme investment strategies become more favorable. When trade-offs are concave, as with specialist trade-offs, only extreme investment strategies are possible. On the other hand, convex trade-offs typically favor intermediate strategies. Depending on the strength of the trade-off, extreme investment may arise through evolutionary branching, in which an evolutionary singularity is convergence stable but not evolutionarily stable [36, 38, 39], For the specialist trade-off analyzed in our model with z = -1, branching was rarely found to occur except at very high temperatures. However, as the strength of the trade-off decreases (z approaches 0), branching can occur more often under moderate conditions (Additional file 1, Section S4). Previous studies have shown that as the strength of a trade-off increases, the evolutionarily singular strategy shifts from an ESS, to a branching point, and finally to a repeller. [37-39]. The weaker, convex trade-offs of generalist mixotrophs on the other hand were found to favor intermediate investment strategies, consistent with true mixotrophic strategies.

The evolutionary outcomes for each trade-off type have implications for the trade-offs likely to be found in extant mixotrophs. Since mixotrophs by definition make use of both photosynthesis and phagotrophy, any trade-off between the two must be capable of maintaining viability when resources are being dedicated to each strategy. For constitutive mixotrophs, the tendency of convex trade-offs to favor intermediate investment strategies over evolutionary time suggests that these trade-offs are likely to be common in mixotrophs. On the other hand, since specialist trade-offs can lead only to pure resource specialization, mixotrophs are not expected to utilize specialist trade-offs, at least over evolutionary timescales sufficiently long to reach an ESS. While mixotrophs with linear trade-offs were found to have intermediate evolutionarily stable investment strategies under a limited range of conditions, they tended to converge towards pure specialization, suggesting that mixotrophs with linear trade-offs can exist, but are likely highly sensitive to environmental parameters. It is also worth noting that even though linear trade-offs are theoretically possible, perfectly linear trade-offs are likely rare in nature [37, 58]. Metabolic specialization arising from mixotrophs, as observed in our model most often for mixotrophs with linear and concave trade-offs, is supported theoretically and observationally [26, 29-30, 59], and long-term temperature shifts which could potentially promote such specialization are known to have taken place throughout Earth’s history [60]

## CONCLUSIONS

In our model, mixotrophs typically responded to increasing temperatures by evolving greater investment in phagotrophy at the expense of photosynthesis. These results suggest that evolution will lead mixotrophs to contribute increasingly to atmospheric CO_2_ flux if ocean temperatures continue to warm several or more degrees as predicted by climate models. However, our results are sensitive to ecological feedback on prey density as well as the intrinsic trade-off between phagotrophy and photosynthesis experienced by mixotrophs. Our results highlight the importance of taking evolution into account in addition to phenotypic plasticity for predicting organismal response to climate change. Although our model focused on the metabolic response of mixotrophs to increasing temperatures, the effects of climate change span many dimensions. Future work should take into account other factors including changes in ocean pH, pCO_2_ and nutrient concentrations and how these factors will synergistically affect aspects of mixotrophic physiology including cell size and prey preference. Theoretical studies predict that mixotrophs have an important effect on the marine carbon cycle [61], but more field and experimental studies will be required to provide support for these predictions. To better understand the influence of mixotrophs on global biogeochemistry and how it will change as they evolve, it will be necessary to better characterize the abundance and metabolic heterogeneity of mixotrophic phytoplankton across different regions of the ocean.

## DECLARATIONS

### Ethics approval and consent to participate

No ethical guidance or approval was required for this study. All methods were carried out in accordance with relevant guidelines and regulations.

### Consent for publication

Not applicable.

### Availability of data and materials

The datasets generated and/or analyzed during the current study are available in the GitHub repository, https://github.com/lg11235/mixotroph-adaptive-dynamics

### Author’s contributions

LMG designed the research, with input from SRP and HVM. LMG and SRP conducted the research, and all authors wrote and revised the paper.

## Acknowledgements

We thank one anonymous reviewer and G.D. Ferriera for comments on an earlier draft of the manuscript.

## REFERENCES

1. IPCC, 2014: Climate Change 2014: Synthesis Report. Contribution of Working Groups I, II and III to the Fifth Assessment Report of the Intergovernmental Panel on Climate Change [Core Writing Team, R.K. Pachauri and L.A. Meyer (eds.)]. IPCC, Geneva, Switzerland, 151 pp.

2. Arrhenius, S. (1889). Über die Dissociationswärme und den Einfluss der Temperatur auf den Dissociationsgrad der Elektrolyte. Zeitschrift Für Physikalische Chemie, 4(1), 96–116.

3. Gillooly, J. F., Brown, J. H., West, G. B., Savage, V. M., & Charnov, E. L. (2001). Effects of size and temperature on metabolic rate. Science, 293(5538), 2248–2251.

4. Doney, S. C., Ruckelshaus, M., Duffy, J. E., Barry, J. P., Chan, F., English, C. A., Galindo, H. M., Grebmeier, J. M., Hollowed, A. B., & Knowlton, N. (2011). Climate change impacts on marine ecosystems.

5. Allen, A. P., Gillooly, J. F., & Brown, J. H. (2005). Linking the global carbon cycle to individual metabolism. Functional Ecology, 19(2), 202–213.

6. Brown, J. H., Gillooly, J. F., Allen, A. P., Savage, V. M., & West, G. B. (2004). TOWARD A METABOLIC THEORY OF ECOLOGY. Ecology, 85(7), 1771–1789. https://doi.org/10.1890/03-9000

7. Yvon-Durocher, G., Allen, A. P., Montoya, J. M., Trimmer, M., & Woodward, G. (2010). Yv. In Advances in Ecological Research (Vol. 43, pp. 267–313). Elsevier. https://doi.org/10.1016/B978-0-12-385005-8.00007-1

8. Monroe, J. G., Markman, D. W., Beck, W. S., Felton, A. J., Vahsen, M. L., & Pressler, Y. (2018). Ecoevolutionary Dynamics of Carbon Cycling in the Anthropocene. Trends in Ecology & Evolution, 33(3), 213–225. https://doi.org/10.1016/j.tree.2017.12.006

9. Field, C. B., Behrenfeld, M. J., Randerson, J. T., & Falkowski, P. (1998). Primary Production of the Biosphere: Integrating Terrestrial and Oceanic Components. Science, 281(5374), 237– 240. https://doi.org/10.1126/science.281.5374.237

10. Thomas, M. K., Kremer, C. T., Klausmeier, C. A., & Litchman, E. (2012). A global pattern of thermal adaptation in marine phytoplankton. Science, 338(6110), 1085–1088.

11. Barton, S., Jenkins, J., Buckling, A., Schaum, C.-E., Smirnoff, N., Raven, J. A., & Yvon-Durocher, G. (2020). Evolutionary temperature compensation of carbon fixation in marine phytoplankton. Ecology Letters, 23(4), 722–733. https://doi.org/10.1111/ele.13469

12. Worden, A. Z., Follows, M. J., Giovannoni, S. J., Wilken, S., Zimmerman, A. E., & Keeling, P. J. (2015). Rethinking the marine carbon cycle: Factoring in the multifarious lifestyles of microbes. Science, 347(6223), 1257594. https://doi.org/10.1126/science.1257594

13. Hartmann, M., Grob, C., Tarran, G. A., Martin, A. P., Burkill, P. H., Scanlan, D. J., & Zubkov, M. V. (2012). Mixotrophic basis of Atlantic oligotrophic ecosystems. Proceedings of the National Academy of Sciences, 109(15), 5756–5760.

14. Ward, B. A., & Follows, M. J. (2016). Marine mixotrophy increases trophic transfer efficiency, mean organism size, and vertical carbon flux. Proceedings of the National Academy of Sciences, 113(11), 2958–2963.

15. Mitra, A., Flynn, K. J., Tillmann, U., Raven, J. A., Caron, D., Stoecker, D. K., Not, F., Hansen, P. J., Hallegraeff, G., Sanders, R., Wilken, S., McManus, G., Johnson, M., Pitta, P., Våge, S., Berge, T., Calbet, A., Thingstad, F., Jeong, H. J., … Lundgren, V. (2016). Defining Planktonic Protist Functional Groups on Mechanisms for Energy and Nutrient Acquisition: Incorporation of Diverse Mixotrophic Strategies. Protist, 167(2), 106–120. https://doi.org/10.1016/j.protis.2016.01.003

16. Polovina, J. J., Howell, E. A., & Abecassis, M. (2008). Ocean’s least productive waters are expanding. Geophysical Research Letters, 35(3), L03618. https://doi.org/10.1029/2007GL031745

17. Stoecker, D. K. (1998). Conceptual models of mixotrophy in planktonic protists and some ecological and evolutionary implications. European Journal of Protistology, 34(3), 281–290. https://doi.org/10.1016/S0932-4739(98)80055-2

18. Flynn KJ, Mitra A, Anestis K, Anschütz AA, Calbet A, Ferreira GD, Gypens N, Hansen PJ, John U, Martin JL, Mansour JS. Mixotrophic protists and a new paradigm for marine ecology: where does plankton research go now?. Journal of Plankton Research. 2019 Jul 26;41(4):375-

19. Wilken, S., Huisman, J., Naus-Wiezer, S., & Van Donk, E. (2013). Mixotrophic organisms become more heterotrophic with rising temperature. Ecology Letters, 16(2), 225–233.

20. Cabrerizo MJ, González-Olalla JM, Hinojosa-López VJ, Peralta-Cornejo FJ, Carrillo P. A shifting balance: responses of mixotrophic marine algae to cooling and warming under UVR. New Phytologist. 2019 Feb;221(3):1317–27.

21. González-Olalla Jm, Medina-Sánchez JM, Carrillo P. Mixotrophic trade-off under warming and UVR in a marine and a freshwater alga. Journal of phycology. 2019 Oct;55(5):1028–40.

22. Princiotta, S. D., Smith, B. T., & Sanders, R. W. (2016). Temperature-dependent phagotrophy and phototrophy in a mixotrophic chrysophyte. Journal of Phycology, 52(3), 432–440.

23. Ferreira GD, Grigoropoulou A, Saiz E, Calbet A. The effect of short-term temperature exposure on vital physiological processes of mixoplankton and protozooplankton. Marine Environmental Research. 2022 Jul 1;179:105693.

24. Flynn, K. J., & Mitra, A. (2009). Building the “perfect beast”: Modelling mixotrophic plankton. Journal of Plankton Research, 31(9), 965–992.

25. Ward, B. A., Dutkiewicz, S., Barton, A. D., & Follows, M. J. (2011). Biophysical aspects of resource acquisition and competition in algal mixotrophs. The American Naturalist, 178(1), 98–112.

26. Raven, J. A. (1997). Phagotrophy in phototrophs. Limnology and Oceanography, 42(1), 198– 205.

27. Hendry, A. (2016). Eco-evolutionary Dynamics. Princeton: Princeton University Press. https://doi.org/10.1515/9781400883080

28. Pelletier, F., Garant, D., & Hendry, A. P. (2009). Eco-evolutionary dynamics. Philosophical Transactions of the Royal Society B: Biological Sciences, 364(1523), 1483–1489. https://doi.org/10.1098/rstb.2009.0027

29. de Castro, F., Gaedke, U., & Boenigk, J. (2009). Reverse Evolution: Driving Forces Behind the Loss of Acquired Photosynthetic Traits. PLoS ONE, 4(12), e8465. https://doi.org/10.1371/journal.pone.0008465

30. Troost, T. A., Kooi, B. W., & Kooijman, S. A. (2005). When do mixotrophs specialize? Adaptive dynamics theory applied to a dynamic energy budget model. Mathematical Biosciences, 193(2), 159–182.

31. Brännström, Å., Johansson, J., & von Festenberg, N. (2013). The Hitchhiker’s Guide to Adaptive Dynamics. Games, 4(3), 304–328. https://doi.org/10.3390/g4030304

32. Diekmann, O. (2003). A beginner’s guide to adaptive dynamics. Mathematical Modelling of Population Dynamics, 47–86. https://doi.org/10.4064/bc63-0-2

33. Levins R. Theory of fitness in a heterogeneous environment. I. The fitness set and adaptive function. The American Naturalist. 1962 Nov 1;96(891):361–73.

34. Lawlor LR, Smith JM. The coevolution and stability of competing species. The American Naturalist. 1976 Jan 1;110(971):79–99.

35. Tilman D. Resources: a graphical-mechanistic approach to competition and predation. The American Naturalist. 1980 Sep 1;116(3):362–93.

36. Schreiber SJ, Tobiason GA. The evolution of resource use. Journal of mathematical biology. 2003 Jul;47(1):56–78.

37. Wickman J, Diehl S, Brännström Å. Evolution of resource specialisation in competitive metacommunities. Ecology letters. 2019 Nov;22(11):1746–56.

38. de Mazancourt C, Dieckmann U. Trade-off geometries and frequency-dependent selection. The American Naturalist. 2004 Dec;164(6):765–78.

39. Rueffler C, Van Dooren TJ, Metz JA. The evolution of resource specialization through frequency-dependent and frequency-independent mechanisms. The American Naturalist. 2006 Jan;167(1):81–93.

40. Huisman, J., & Weissing, F. J. (1994). Light-limited growth and competition for light in well-mixed aquatic environments: An elementary model. Ecology, 75(2), 507–520.

41. Moeller, H. V., Neubert, M. G., & Johnson, M. D. (2019). Intraguild predation enables coexistence of competing phytoplankton in a well-mixed water column. Ecology, 100(12). https://doi.org/10.1002/ecy.2874

42. Holling, C. S. (1965). The functional response of predators to prey density and its role in mimicry and population regulation. The Memoirs of the Entomological Society of Canada, 97(S45), 5–60.

43. Vasseur, D. A., & McCann, K. S. (2005). A Mechanistic Approach for Modeling Temperature-Dependent Consumer-Resource Dynamics. The American Naturalist, 166(2), 184–198. https://doi.org/10.1086/431285

44. Dell, A. I., Pawar, S., & Savage, V. M. (2011). Systematic variation in the temperature dependence of physiological and ecological traits. Proceedings of the National Academy of Sciences, 108(26), 10591–10596. https://doi.org/10.1073/pnas.1015178108

45. Metz, J.A., Geritz, S.A., Meszéna, G., Jacobs, F.J., & Heerwaarden, J.V. (1995). Adaptive Dynamics: A Geometrical Study of the Consequences of Nearly Faithful Reproduction.

46. Dieckmann, U., & Law, R. (1996). The dynamical theory of coevolution: a derivation from stochastic ecological processes. Journal of mathematical biology, 34(5), 579–612.

47. Geritz, S. A. H., Kisdi, É., Meszéna, G., & Metz, J. A. J. (1998). Evolutionarily singular strategies and the adaptive growth and branching of the evolutionary tree. Evolutionary Ecology, 12(1), 35–57. https://doi.org/10.1023/A:1006554906681

48. Geritz, S. A. H. (2005). Resident-invader dynamics and the coexistence of similar strategies. Journal of Mathematical Biology, 50(1), 67–82. https://doi.org/10.1007/s00285-004-0280-8

49. Dercole, F., & Geritz, S. A. H. (2016). Unfolding the resident–invader dynamics of similar strategies. Journal of Theoretical Biology, 394, 231–254. https://doi.org/10.1016/j.jtbi.2015.11.032

50. Lion, S. (2018). Theoretical Approaches in Evolutionary Ecology: Environmental Feedback as a Unifying Perspective. The American Naturalist, 191(1), 21–44. https://doi.org/10.1086/694865

51. Collins, S., & De Meaux, J. (2009). Adaptation to different rates of environmental change in Chlamydomonas. Evolution: International Journal of Organic Evolution, 63(11), 2952–2965.

52. Lepori-Bui M, Paight C, Eberhard E, Mertz C, Moeller HV. (in press). Evidence for evolutionary adaptation of mixotrophic nanoflagellates to warmer temperatures. Global Change Biology.

53. Lenski RE, Bennett AF. Evolutionary response of Escherichia coli to thermal stress. The American Naturalist. 1993 Jul 1;142:S47–64.

54. Kaitala V, Hiltunen T, Becks L, Scheuerl T. Co-evolution as an important component explaining microbial predator-prey interaction. Journal of Theoretical Biology. 2020 Feb 7;486:110095.

55. Eppley, R. W., & Peterson, B. J. (1979). Particulate organic matter flux and planktonic new production in the deep ocean. Nature, 282(5740), 677–680. https://doi.org/10.1038/282677a0

56. Hallsson LR, Björklund M. Selection in a fluctuating environment leads to decreased genetic variation and facilitates the evolution of phenotypic plasticity. Journal of evolutionary biology. 2012 Jul;25(7):1275–90.

57. Lande R. Evolution of phenotypic plasticity and environmental tolerance of a labile quantitative character in a fluctuating environment. Journal of evolutionary biology. 2014 May;27(5):866–75.

58. Caetano, R. A., Ispolatov, Y., & Doebeli, M. (2020). Evolution of diversity in metabolic strategies [Preprint]. Evolutionary Biology. https://doi.org/10.1101/2020.10.20.347419

59. Fawcett, R. C., & Parrow, M. W. (2014). Mixotrophy and loss of phototrophy among geographic isolates of freshwater Esoptrodinium / Bernardinium sp. (Dinophyceae). Journal of Phycology, 50(1), 55–70. https://doi.org/10.1111/jpy.12144

60. Milankovitch, M. (1920). Mathematical Theory of Heat Phenomena Produced by Solar Radiation. Gauthier-Villars.

61. Ward BA, Follows MJ. Marine mixotrophy increases trophic transfer efficiency, mean organism size, and vertical carbon flux. Proceedings of the National Academy of Sciences. 2016 Mar 15;113(11):2958–63.

