## Additional File 1 for "Modeling the metabolic evolution of mixotrophic phytoplankton in response to rising ocean surface temperatures"

Name: Additional file 1

Format: .pdf

Title of data: SUPPLEMENTARY MATERIAL: MODELING THE METABOLIC EVOLUTION OF MIXOTROPHIC PHYTOPLANKTON IN RESPONSE TO RISING OCEAN SURFACE TEMPERATURES

Description: Supplementary figures and analytical work to support manuscript results.

### SUPPLEMENTARY MATERIAL:

### MODELING THE METABOLIC EVOLUTION OF MIXOTROPHIC PHYTOPLANKTON IN RESPONSE TO RISING OCEAN SURFACE TEMPERATURES

Logan M. Gonzalez, Stephen R. Proulx, and Holly V. Moeller

September 2022

#### S1 Supplementary Figures

To test the effects of our assumption of linear thermal responses on our model's predictions, we performed additional analyses in which photosynthesis and phagotrophy are modeled as exponential functions of temperature.

We used the Arrhenius equation (Arrhenius 1889) to determine k, the rate constant:

$$\mathrm{k} = \mathrm{A}e^{\frac{-\mathrm{E_a}}{\mathrm{k_B T}}} \tag{S1}$$

where T is the absolute temperature in Kelvin, $\mathrm{k_B}$ is the Boltzmann constant ($8.617 \cdot 10^{-5}$ eV $\mathrm{K}^{-1}$), A is the normalization constant, and the activation energies $\mathrm{E_a}$ are 0.50 eV for photosynthesis and 0.85 eV for phagotrophy (from Wilken et al., 2013). Normalization constants for phagotrophy and photosynthesis ($\mathrm{A}_\alpha$ and $\mathrm{A}_\rho$, respectively) were chosen such that at the baseline temperature (286 K), k = 1:

$$\mathrm{A}_\alpha = \frac{1}{e^{\frac{-\mathrm{E_a}}{\mathrm{k_B T}}}} \approx 9.413 \cdot 10^{14} \tag{S2}$$

$$A_\rho = \frac{1}{e^{\frac{-\mathrm{E_a}}{\mathrm{k_B T}}}} \approx 6.428 \cdot 10^{8} \tag{S3}$$

We multiplied maximum carbon uptake rate, $\rho_{\max}$, and maximum attack rate of mixotroph on bacteria,

$\alpha_{\max}$ by their respective k values to determine the temperature dependent rates $\rho_{\max}(\mathrm{T})$ and $\alpha_{\max}(\mathrm{T})$. Results of the evolutionary analysis are shown in Figure S1.

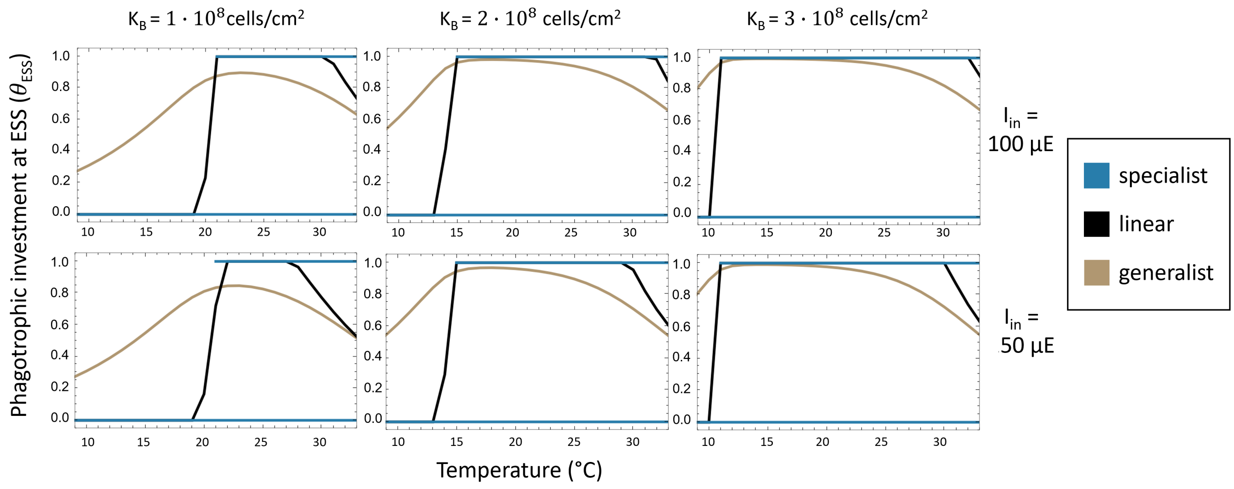


Figure S1: $\theta_{ESS}$ as a function of temperature, light, and prey carrying capacity for mixotrophs with specialist (blue), generalist (light brown), and linear (black) trade-offs. Phagotrophy and photosynthesis are modeled as exponential functions of temperature, according to the Arrhenius equation (Arrhenius 1889), with $\mathrm{k_B} = 8.617 \cdot 10^{-5}$ eV $\mathrm{K}^{-1}$ Activation energies measured for carbon fixation and grazing from Wilken et al. 2013 (0.50 eV, 0.85 eV, respectively) were used to parameterize functions. Normalization constants of $9.413 \cdot 10^{14}$ and $6.428 \cdot 10^{8}$ were chosen such that at the baseline temperature, $\mathrm{T_0}$, $\rho_{\max}$ and $\alpha_{\max}$ were scaled by 1. Specialist mixotrophs exhibited evolutionary bistability at $\theta_{\mathrm{ESS}}$=0 and $\theta_{\mathrm{ESS}}$=1, however mixotroph population sizes were negative in some cases for $\theta_{\mathrm{ESS}} = 1$ at low temperatures.

#### S2 Model overview

Our model for the population dynamics of the interacting mixotroph and bacteria prey is

$$\frac{\mathrm{d}M}{\mathrm{d}t} = \left[\frac{\rho(\theta,\mathrm{z},\mathrm{T})}{\mathrm{k}M}\ln\left(\frac{\mathrm{h}+\mathrm{I_{in}}}{\mathrm{h}+\mathrm{I_{in}}e^{-\mathrm{k}M}}\right)+\mathrm{b}\alpha(\theta,\mathrm{T})B-\mathrm{l}\right]M \tag{S4}$$

$$\frac{\mathrm{d}B}{\mathrm{d}t} = \left[\mathrm{r}(1-B/\mathrm{K_B})-\alpha(\theta,\mathrm{T})M\right]B \tag{S5}$$

Note that the density of the mixotrophs enters into the calculation for the amount of light available for photosynthesis.

We further define

$$\rho(\theta,\mathrm{z},\mathrm{T}) = \rho_{\max}(1-\theta^{2^{\mathrm{z}}})^{\frac{1}{2^{\mathrm{z}}}}(\max(0,\mathrm{m}_\rho(\mathrm{T}-\mathrm{T}_{0\rho}))) \tag{S6}$$

$$\alpha(\theta,\mathrm{T}) = \alpha_{\max}\theta(\max(0,\mathrm{m}_\alpha(\mathrm{T}-\mathrm{T}_{0\alpha}))) \tag{S7}$$

In order to determine whether or not a mutant mixotroph strategy replaces a resident strategy we must define the invasion fitness of the mutant. We note that for the resident strategy $\theta_{\mathrm{res}}$, the equilibrium values for equations S4 and S5 are denoted $M^*(\theta_{\mathrm{res}})$ and $B^*(\theta_{\mathrm{res}})$.

The differential equation describing mutant mixotroph population growth rate, assuming that the mutant is sufficiently rare that the resident's phenotype determines both the amount of light available and the population dynamics of the bacteria, is

$$\frac{\mathrm{d}M_{\mathrm{mut}}}{\mathrm{d}t} = \left[\frac{\rho(\theta_{\mathrm{mut}},\mathrm{z},\mathrm{T})}{kM^*(\theta_{\mathrm{res}})}\ln\left(\frac{\mathrm{h}+\mathrm{I_{in}}}{\mathrm{h}+\mathrm{I_{in}}e^{-\mathrm{k}M^*(\theta_{\mathrm{res}})}}\right)+\mathrm{b}\alpha(\theta_{\mathrm{mut}},\mathrm{T})B^*(\theta_{\mathrm{res}})-\mathrm{l}\right]M_{\mathrm{mut}} \tag{S8}$$

This is a linear equation in $M_{\mathrm{mut}}$, therefore we can define the invasion exponent for the mutant population as

$$\lambda(\theta_{\mathrm{mut}},\theta_{\mathrm{res}}) = \frac{1}{M_{\mathrm{mut}}}\frac{\mathrm{d}M_{\mathrm{mut}}}{\mathrm{d}t} \tag{S9}$$

When $\lambda(\theta_{\mathrm{mut}},\theta_{\mathrm{res}}) > 0$, the mutant is able to invade and we assume that it then replaces the resident mixotroph strategy. This invasion-replacement process continues until the population reaches an evolutionarily stable state, ESS, at which point the system is locally stable to further mutant invasion.

In order to find the ESS we calculate the selection gradient (Brännström et al. 2013) by differentiating

$\lambda(\theta_{\mathrm{mut}}, \theta_{\mathrm{res}})$ with respect to $\theta_{\mathrm{mut}}$ and setting $\theta_{\mathrm{mut}} = \theta_{\mathrm{res}}$.

$$\begin{aligned}
\left.\frac{\mathrm{d}\lambda(\theta_{\mathrm{mut}}, \theta_{\mathrm{res}})}{\mathrm{d}\theta_{\mathrm{mut}}}\right|_{\theta_{\mathrm{mut}}=\theta_{\mathrm{res}}} &= \frac{\mathrm{d}\rho(\theta_{\mathrm{res}}, \mathrm{z}, \mathrm{T})}{\mathrm{d}\theta_{\mathrm{res}}} \frac{1}{\mathrm{k}M^*(\theta_{\mathrm{res}})} \ln\left(\frac{\mathrm{h} + \mathrm{I}_{\mathrm{in}}}{\mathrm{h} + \mathrm{I}_{\mathrm{in}} e^{-\mathrm{k}M^*(\theta_{\mathrm{res}})}}\right) + \mathrm{b}\frac{\mathrm{d}\alpha(\theta_{\mathrm{res}}, \mathrm{T})}{\mathrm{d}\theta_{\mathrm{res}}} B^*(\theta_{\mathrm{res}}) \\
&= b\alpha(T)B^*(\theta_{\mathrm{res}}) - \frac{\rho(\theta_{\mathrm{res}}, \mathrm{T})\theta_{\mathrm{res}}^{2^{\mathrm{z}}-1}}{\mathrm{k}(1 - \theta_{\mathrm{res}}^{2^{\mathrm{z}}})M^*(\theta_{\mathrm{res}})} \ln\left(\frac{\mathrm{h} + \mathrm{I}}{\mathrm{h} + \mathrm{I}e^{-\mathrm{k}M^*(\theta_{\mathrm{res}})}}\right) \qquad \text{(S10)}
\end{aligned}$$

The selection gradient allows us to determine the evolutionarily stable phagotrophic investment strategy, $\theta_{\mathrm{ESS}}(\mathrm{T})$, that a population of mixotrophs is predicted to evolve towards at each given temperature. $\theta_{\mathrm{ESS}}(\mathrm{T})$ is found by calculating $\theta_{\mathrm{res}}$ when $\frac{\mathrm{d}\lambda(\theta_{\mathrm{mut}},\theta_{\mathrm{res}})}{\mathrm{d}\theta_{\mathrm{mut}}}|_{\theta_{\mathrm{mut}}=\theta_{\mathrm{res}}} = 0$ and $\frac{\mathrm{d}^2\lambda(\theta_{\mathrm{mut}},\theta_{\mathrm{res}})}{\mathrm{d}\theta_{\mathrm{mut}}^2} < 0$. When these conditions are met, $\theta_{\mathrm{res}}$ represents a fitness maximum defined as the evolutionarily stable state, $\theta_{\mathrm{ESS}}$. If $\frac{\mathrm{d}^2\lambda(\theta_{\mathrm{mut}},\theta_{\mathrm{res}})}{\mathrm{d}\theta_{\mathrm{mut}}^2}|_{\theta_{\mathrm{mut}}=\theta_{\mathrm{res}}} > 0$, $\theta_{\mathrm{res}}$ represents a fitness minimum that will be either a repeller or branching point. If it is a branching point, the strategy of the population will converge towards the evolutionarily singular strategy before splitting to become dimorphic. Since there are no closed form solutions for $M^*$ and $B^*$, $\theta_{\mathrm{ESS}}(\mathrm{T})$ was solved numerically.

#### S3 Evolutionary stable states are not yield-maximizing

In this section we show that, when mixotrophs evolve to an un-invadable strategy (ESS), their population densities are lower than could be achieved with other investment strategies. We show this by finding implicit derivatives of the equilibrium population size and showing that, at the ESS, mixotroph equilibrium density cannot exist at a population maximum. This indicates that evolution does not maximize mixotroph population size.

The ESS condition from equation S10 gives

$$\frac{\mathrm{d}\rho(\theta_{\mathrm{res}}, \mathrm{z}, \mathrm{T})}{\mathrm{d}\theta_{\mathrm{res}}} \frac{1}{\mathrm{k}M^*(\theta_{\mathrm{res}})} \ln\left(\frac{\mathrm{h}+\mathrm{I}_{\mathrm{in}}}{\mathrm{h}+\mathrm{I}_{\mathrm{in}}e^{-\mathrm{k}M^*(\theta_{\mathrm{res}})}}\right) + b\frac{\mathrm{d}\alpha(\theta_{\mathrm{res}}, T)}{\mathrm{d}\theta_{\mathrm{res}}} B^*(\theta_{\mathrm{res}}) = 0 \tag{S11}$$

We will use this condition to draw conclusions about how equilibrium mixotroph density changes with mixotroph strategy.

The population densities at equilibrium can be found by setting equations S4 and S5 equal to 0 and finding the non-trivial values $M^*$ and $B^*$ that solve these equalities.

$$0 = \left[\frac{\rho(\theta, \mathrm{z}, \mathrm{T})}{\mathrm{k}M^*} \ln\left(\frac{\mathrm{h}+\mathrm{I}_{\mathrm{in}}}{\mathrm{h}+\mathrm{I}_{\mathrm{in}}e^{-\mathrm{k}M^*}}\right) + \mathrm{b}\alpha(\theta, \mathrm{T})B^* - 1\right] M^* \tag{S12}$$

$$0 = \left[\mathrm{r}(1 - B^*/\mathrm{K}_{\mathrm{B}}) - \alpha(\theta, \mathrm{T})M^*\right] B^* \tag{S13}$$

Now we note that we can consider the equilibrium values to be functions of $\theta$ and differentiate both equations with respect to $\theta$. For the $M$ equation we get

$$\frac{\mathrm{d}\rho(\theta, \mathrm{z}, \mathrm{T})}{\mathrm{d}\theta} \frac{1}{\mathrm{k}M^*} \ln\left(\frac{\mathrm{h}+\mathrm{I}_{\mathrm{in}}}{\mathrm{h}+\mathrm{I}_{\mathrm{in}}e^{-\mathrm{k}M^*}}\right) + \rho(\theta, \mathrm{z}, \mathrm{T})\frac{\mathrm{d}}{\mathrm{d}\theta}[\frac{1}{\mathrm{k}M^*} \ln\left(\frac{\mathrm{h}+\mathrm{I}_{\mathrm{in}}}{\mathrm{h}+\mathrm{I}_{\mathrm{in}}e^{-\mathrm{k}M^*}}\right)] + \mathrm{b}\frac{\mathrm{d}\alpha(\theta, \mathrm{T})}{\mathrm{d}\theta} B^* + \mathrm{b}\alpha(\theta, \mathrm{T})\frac{\mathrm{d}B^*}{\mathrm{d}\theta} = 0$$

Substituting in the ESS conditions and noting that $\frac{\mathrm{df}(M(\theta))}{\mathrm{d}\theta} = \frac{\mathrm{df}(M(\theta))}{\mathrm{d}M(\theta)} \frac{\mathrm{d}M(\theta)}{\mathrm{d}\theta}$ we get

$$\rho(\theta, \mathrm{z}, \mathrm{T})\frac{\mathrm{d}}{\mathrm{d}M^*}[\frac{1}{\mathrm{k}M^*} \ln\left(\frac{\mathrm{h}+\mathrm{I}_{\mathrm{in}}}{\mathrm{h}+\mathrm{I}_{\mathrm{in}}e^{-\mathrm{k}M^*}}\right)]\frac{\mathrm{d}M^*}{\mathrm{d}\theta} + \mathrm{b}\alpha(\theta, \mathrm{T})\frac{\mathrm{d}B^*}{\mathrm{d}\theta} = 0 \tag{S14}$$

The implication of equation S14 is that if the ESS value of $\theta$ maximizes mixotroph density, both $\frac{\mathrm{d}M^*}{\mathrm{d}\theta} = 0$ and $\frac{\mathrm{d}B^*}{\mathrm{d}\theta} = 0$. I.E. if $M^*$ is at a local extremum then so is $B^*$. Turning back to the $B$ equation we take the implicit derivative with respect to $\theta$ and get

$$\frac{\mathrm{d}\alpha(\theta,\mathrm{T})}{\mathrm{d}\theta}M^* = \frac{-\mathrm{r}\frac{\mathrm{d}B^*}{\mathrm{d}\theta}}{\mathrm{K_B}} - \alpha(\theta,\mathrm{T})\frac{\mathrm{d}M^*}{\mathrm{d}\theta} \tag{S15}$$

Substituting in $\frac{\mathrm{d}M^*}{\mathrm{d}\theta} = 0$ and $\frac{\mathrm{d}B^*}{\mathrm{d}\theta} = 0$ we see that this would imply that

$$M^* = 0. \tag{S16}$$

Thus, the ESS value of $\theta$ could only result in $M^*$ being a local maximum if $M^* = 0$, which cannot be true for any situation where the mixotroph is not extinct. We conclude that the evolutionary-ecology process does not maximize mixotroph population density.

### S4 Specialist ESS and branching conditions

In this section, we show that for specialist mixotrophs, there exist evolutionary critical points near $\theta = 0$ and $\theta = 1$ that are always repeller points, meaning they are neither convergence stable nor evololutionarily stable. We show this by analyzing the fitness gradient for a specialist mixotrophs and showing that constraints on the values at the boundaries $\theta = 0$ and $\theta = 1$ result in the presence of at least one repeller point. We also show that specialists are capable of satisfying the conditions for evolutionary branching, indicating the existence of an additional evolutionarily critical point in some cases.

Let $s(\theta_{\rm res}) = \frac{{\rm d}\lambda(\theta_{\rm mut},\theta_{\rm res})}{{\rm d}\theta_{\rm mut}}|_{(\theta_{\rm mut}=\theta_{\rm res})}$. For a population of mixotrophs at constant temperature,

$$\begin{aligned} {\rm s}(\theta_{\rm res}) &= {\rm b}\alpha_{\rm max}B^*(\theta_{\rm res}) - \\ &\left(\frac{\rho_{\rm max}}{{\rm k}M^*(\theta_{\rm res})}\ln\left(\frac{{\rm h}+{\rm I_{in}}}{{\rm h}+{\rm I_{in}}e^{-{\rm k}M^*(\theta_{\rm res})}}\right)\right)\theta_{\rm res}^{(-1+2^{\rm z})}(1-\theta_{\rm res}^{2^{\rm z}})^{(-1+2^{-\rm z})} \end{aligned} \quad \text{(S17)}$$

Assume that $B^*(\theta) > 0$, $M^*(\theta) > 0$. Note that when $z < 0$, ${\rm s}(\theta)$ approaches $-\infty$ as $\theta$ approaches 0, and that the value of s(1) is greater than 0. Because s is a continuous and differentiable function, application of the intermediate value theorem shows that ${\rm s}(\theta) = 0$ for some values of $\theta$. Call $\theta_{\rm min}$ the smallest value of $\theta$ such that $s(\theta_{\rm min}) = 0$. Because s is continuous, and takes on a negative value at $0 < \theta_{\rm min}$, we have ${\rm s}'(\theta_{\rm min}) \geq 0$. Likewise, call $\theta_{\rm max}$ the largest value of $\theta$ such that $s(\theta_{\rm max}) = 0$. Because s is continuous, and takes on a positive value at $1 > \theta_{\rm max}$, we have ${\rm s}'(\theta_{\rm max}) \geq 0$.

Therefore near either 0 or 1, the nearest evolutionary critical point is a repeller, indicating that extreme values of $\theta$ are evolutionarily stable.

In order for evolutionary branching to be possible, two conditions must be met at a third evolutionary critical point in between the repellers:

$$\frac{{\rm d}^2\lambda(\theta_{\rm mut},\theta_{\rm res})}{{\rm d}\theta_{\rm mut}^2} > 0 \quad \text{(S18)}$$

$$\frac{{\rm d}s(\theta_{\rm res})}{{\rm d}\theta_{\rm res}} < 0 \quad \text{(S19)}$$

For specialist trade-off mixotrophs, the first condition is always satisfied, assuming $M^*(\theta_{\rm res}) \geq 0$ and $0 \leq \theta_{\rm res} \leq 1$,

$$\frac{{\rm d}^2\lambda(\theta_{\rm mut},\theta_{\rm res})}{{\rm d}\theta_{\rm mut}^2} = \frac{\rho(1-2^{\rm z})(\theta_{\rm mut}^{(-2+2^{\rm z})})(1-\theta_{\rm mut}^{2^{\rm z}})^{(-2+2^{-\rm z})}\ln\left(\frac{{\rm h}+{\rm I_{in}}}{{\rm h}+{\rm I_{in}}e^{-{\rm k}M^*(\theta_{\rm res}))}}\right)}{{\rm k}M^*(\theta_{\rm res})} > 0 \quad \text{(S20)}$$

For the second branching condition to be satisfied, the following must be true at an evolutionarily singular strategy

$$\frac{\mathrm{d}}{\mathrm{d}\theta_{\mathrm{res}}}\left[\frac{\rho}{\mathrm{k}M^*(\theta_{\mathrm{res}})}\ln\left(\frac{\mathrm{h}+\mathrm{I_{in}}}{\mathrm{h}+\mathrm{I_{in}}e^{-\mathrm{k}M^*(\theta_{\mathrm{res}})}}\right)\theta_{\mathrm{res}}^{(-1+2^{\mathrm{z}})}(1-\theta_{\mathrm{res}}^{2^{\mathrm{z}}})^{(-1+2^{-\mathrm{z}})}\right] > \frac{\mathrm{d}}{\mathrm{d}\theta_{\mathrm{res}}}[\mathrm{b}\alpha B^*(\theta_{\mathrm{res}})] \tag{S21}$$

This can occur if the abundance of bacteria decreases significantly with increasing $\theta$, which is expected to occur at elevated temperatures when baseline grazing rates are large. The trade-off parameter z is also expected to influence conditions for branching. Let $\mathrm{P}(\theta_{\mathrm{res}}) = \frac{\rho}{\mathrm{k}M^*(\theta_{\mathrm{res}})}\ln\left(\frac{\mathrm{h}+\mathrm{I_{in}}}{\mathrm{h}+\mathrm{I_{in}}e^{-\mathrm{k}M^*(\theta_{\mathrm{res}})}}\right)$, $\mathrm{f}(\theta_{\mathrm{res}}) = \theta_{\mathrm{res}}^{(-1+2^{\mathrm{z}})}(1-\theta_{\mathrm{res}}^{2^{\mathrm{z}}})^{(-1+2^{-\mathrm{z}})}$, and $\mathrm{G}(\theta_{\mathrm{res}}) = \mathrm{b}\alpha B^*(\theta_{\mathrm{res}})$. Using the product rule, equation S21 becomes

$$\frac{\mathrm{dP}(\theta_{\mathrm{res}})}{\mathrm{d}\theta_{\mathrm{res}}}\mathrm{f}(\theta_{\mathrm{res}}) + \mathrm{P}(\theta_{\mathrm{res}})\frac{\mathrm{df}(\theta_{\mathrm{res}})}{\mathrm{d}\theta_{\mathrm{res}}} > \frac{\mathrm{dG}(\theta_{\mathrm{res}})}{\mathrm{d}\theta_{\mathrm{res}}} \tag{S22}$$

We can see from figure S2 that $\frac{\mathrm{df}(\theta_{\mathrm{res}})}{\mathrm{d}\theta_{\mathrm{res}}} < 0$, $\lim_{\mathrm{z}\to 0}\frac{\mathrm{df}(\theta_{\mathrm{res}})}{\mathrm{d}\theta_{\mathrm{res}}}|_{\theta_{\mathrm{res}}>0} = 0$, and $\lim_{\mathrm{z}\to 0}\mathrm{f}(\theta_{\mathrm{res}}) = 1$, This implies that $\lim_{\mathrm{z}\to 0}[\frac{\mathrm{dP}(\theta_{\mathrm{res}})}{\mathrm{d}\theta_{\mathrm{res}}}\mathrm{f}(\theta_{\mathrm{res}}) + P(\theta_{\mathrm{res}})\frac{\mathrm{df}(\theta_{\mathrm{res}})}{\mathrm{d}\theta_{\mathrm{res}}}] = \frac{\mathrm{d}P(\theta_{\mathrm{res}})}{\mathrm{d}\theta_{\mathrm{res}}}$.

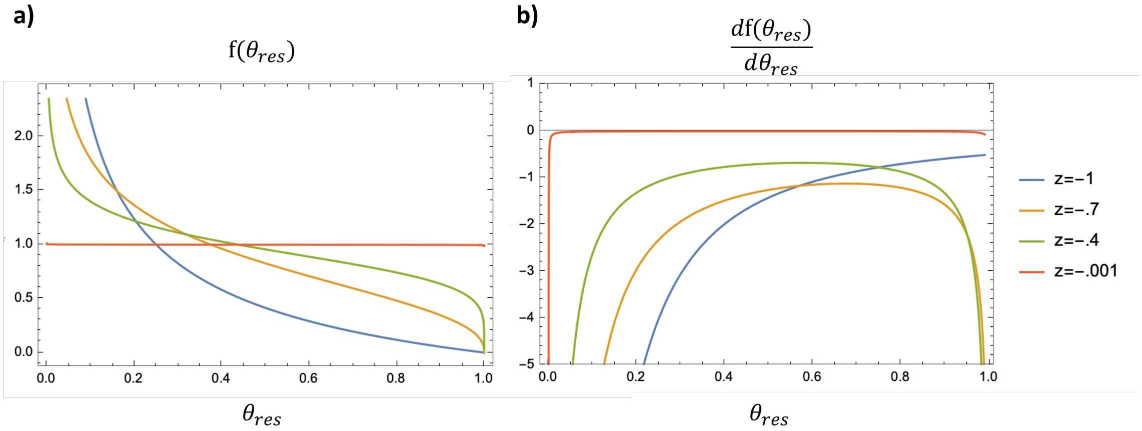


Figure S2: Plots of $\mathrm{f}(\theta_{\mathrm{res}})$ and $\frac{\mathrm{df}(\theta_{\mathrm{res}})}{\mathrm{d}\theta_{\mathrm{res}}}$ vs $\theta_{\mathrm{res}}$ as z is varied. As z approaches 0, $f(\theta_{\mathrm{res}})$ approaches 1 and $\frac{\mathrm{df}(\theta_{\mathrm{res}})}{\mathrm{d}\theta_{\mathrm{res}}}$ approaches 0.

Assuming $M^*(\theta_{\mathrm{res}}) > 0$, $\mathrm{P}(\theta_{\mathrm{res}}) > 0$, then $\mathrm{P}(\theta_{\mathrm{res}})\frac{\mathrm{df}(\theta_{\mathrm{res}})}{\mathrm{d}\theta_{\mathrm{res}}}$ is strictly negative. Therefore, $\frac{\mathrm{dP}(\theta_{\mathrm{res}})}{\mathrm{d}\theta_{\mathrm{res}}} \geq \frac{\mathrm{dP}(\theta_{\mathrm{res}})}{\mathrm{d}\theta_{\mathrm{res}}}\mathrm{f}(\theta_{\mathrm{res}}) + \mathrm{P}(\theta_{\mathrm{res}})\frac{\mathrm{df}(\theta_{\mathrm{res}})}{\mathrm{d}\theta_{\mathrm{res}}}$. Assuming the minimal value of $\frac{\mathrm{dP}(\theta_{\mathrm{res}})}{\mathrm{d}\theta_{\mathrm{res}}}$, this relationship can be used to substitute $\mathrm{P}(\theta_{\mathrm{res}})\frac{\mathrm{df}(\theta_{\mathrm{res}})}{\mathrm{d}\theta_{\mathrm{res}}}$ in equation S22 to obtain the following

$$\frac{\mathrm{dP}(\theta_{\mathrm{res}})}{\mathrm{d}\theta_{\mathrm{res}}}\mathrm{f}(\theta_{\mathrm{res}}) + \frac{\mathrm{dP}(\theta_{\mathrm{res}})}{\mathrm{d}\theta_{\mathrm{res}}}[1-\mathrm{f}(\theta_{\mathrm{res}})] = 0 \tag{S23}$$

Thus, when z is near to zero, the relationship in equation S22 will hold assuming $\frac{\mathrm{dG}(\theta_{\mathrm{res}})}{\mathrm{d}\theta_{\mathrm{res}}} < 0$ and resulting in a branching point. Thus we conclude that at least for small values of z, there is an interior branching point.

#### S5 $\theta_{\mathrm{ESS}}$ at high temperatures

Assuming phagotrophic investment is fixed, prey abundance as a function of temperature at equilibrium is given by equation S24

$$B^* = \mathrm{K_B}(1 - \frac{\alpha(\mathrm{T})M^*}{\mathrm{r}}) \tag{S24}$$

If $\alpha(\mathrm{T})$ is allowed to increase without bound and assuming $M^*$ is positive, there exists a temperature at which $B^* = 0$ and grazing provides no benefit to mixotrophs. Including a phagotrophic investment term $\theta$ that can vary freely from 0 to 1 prevents this from ever happening, as mixotrophs can evolutionarily adjust $\theta$ to always maintain positive values of $B^*$ if $0 \leq \theta \leq 1$ and $0 \leq \alpha(T) < \infty$

$$B^* = \mathrm{K_B}(1 - \frac{\alpha(\mathrm{T})\theta M^*}{\mathrm{r}}) \tag{S25}$$

Thus, as temperature is increased to the point where per-capita grazing rates are too large to be beneficial to the mixotroph population as a whole, phagotrophic investment $\theta$ is expected to decrease.

#### S6 Linear-trade-off mixotroph equilibrium abundance at $\theta_{\mathrm{ESS}}$ is equivalent to equilibrium abundance of strict phototrophs

In this section, we show that a linear trade-off function for mixotrophs results in the equivalence between mixotroph equilibrium abundance at an ESS and equilibrium abundance of strict phototrophs with an investment strategy of $\theta = 0$. We show this by examining the selection gradient and showing that at an ESS, a linear trade-off results in the potential photosynthetic and phagotrophic growth rates (rates of photosynthesis and phagotrophy independent of $\theta$) being equivalent. Using this relationship, the phagotrophic growth rate component of the mixotroph growth rate can be substituted, resulting in an equation that is equivalent to the growth rate of a strict phototroph, indicating that regardless of the value of $\theta_{\mathrm{ESS}}$, equilibrium abundance must be equivalent to equilibrium abundance when $\theta = 0$. We note that this does not apply to the equilibrium abundance of strict heterotrophs since there would still be dependence on the specific value of $\theta_{\mathrm{ESS}}$ through the bacterial prey growth equation.

The temperature-independent selection gradient for a mixotroph is defined as:

$$\left.\frac{\mathrm{d}\lambda(\theta_{\mathrm{mut}},\theta_{\mathrm{res}})}{\mathrm{d}\theta_{\mathrm{mut}}}\right|_{\theta_{\mathrm{mut}}=\theta_{\mathrm{res}}} = \mathrm{b}\alpha B^* - \frac{\rho(\theta_{\mathrm{res}})\theta_{\mathrm{res}}^{2^{\mathrm{z}}-1}}{\mathrm{k}(1-\theta_{\mathrm{res}}^{2^{\mathrm{z}}})M^*}\ln\left(\frac{\mathrm{h}+\mathrm{I_{in}}}{\mathrm{h}+\mathrm{I_{in}}e^{-\mathrm{k}M^*}}\right) \tag{S26}$$

where $\rho(\theta_{\mathrm{res}}) = \rho(1-{\theta_{\mathrm{res}}^2}^{\mathrm{z}})^{\frac{1}{2^{\mathrm{z}}}}$. For linear-trade-off mixotrophs with z = 0, this equation becomes:

$$\left.\frac{\mathrm{d}\lambda(\theta_{\mathrm{mut}},\theta_{\mathrm{res}})}{\mathrm{d}\theta_{\mathrm{mut}}}\right|_{\theta_{\mathrm{mut}}=\theta_{\mathrm{res}}} = \mathrm{b}\alpha B^* - \frac{\rho}{\mathrm{k}M^*}\ln\left(\frac{\mathrm{h}+\mathrm{I_{in}}}{\mathrm{h}+\mathrm{I_{in}}e^{-\mathrm{k}M^*}}\right) \tag{S27}$$

The selection gradient must equal 0 at an ESS, leading to the following:

$$\mathrm{b}\alpha B^* = \frac{\rho}{\mathrm{k}M^*}\ln\left(\frac{\mathrm{h}+\mathrm{I_{in}}}{\mathrm{h}+\mathrm{I_{in}}e^{-\mathrm{k}M^*}}\right) \tag{S28}$$

This can be used to substitute the phagotrophic component of the mixotroph growth rate for linear-trade-off mixotrophs:

$$\begin{aligned}\frac{1}{M^*}\frac{\mathrm{d}M}{\mathrm{d}t} &= \theta\frac{\rho}{\mathrm{k}M^*}\ln\left(\frac{\mathrm{h}+\mathrm{I_{in}}}{\mathrm{h}+\mathrm{I_{in}}e^{-\mathrm{k}M^*}}\right) + (1-\theta)\frac{\rho}{\mathrm{k}M^*}\ln\left(\frac{\mathrm{h}+\mathrm{I_{in}}}{\mathrm{h}+\mathrm{I_{in}}e^{-\mathrm{k}M^*}}\right) - l \\ &= \frac{\rho}{\mathrm{k}M^*}\ln\left(\frac{\mathrm{h}+\mathrm{I_{in}}}{\mathrm{h}+\mathrm{I_{in}}e^{-\mathrm{k}M^*}}\right) - \mathrm{l} \\ &= 0\end{aligned} \tag{S29}$$

In the above equation, $\frac{\rho}{\mathrm{k}M^*}\ln\left(\frac{\mathrm{h}+\mathrm{I_{in}}}{\mathrm{h}+\mathrm{I_{in}}e^{-\mathrm{k}M^*}}\right)-\mathrm{l}$ is the growth rate for strict phototrophs at equilibrium. Therefore at an ESS, the growth rate equation for linear-trade-off mixotrophs is equivalent to that of strict phototrophs and will result in these two populations having the same equilibrium abundances. This is a direct consequence of a linear trade-off function and nonlinear trade-offs do not satisfy these conditions.
